# Development of CRISPR/Cas13-based analytical tools to study RNA-Protein Interactions

**DOI:** 10.1101/2025.04.22.649962

**Authors:** Keita Yamamoto, Shiori Shikata, Yangying Hao, Takuya Tomita, Els Verhoeyen, Yasushi Saeki, Susumu Goyama

**Affiliations:** Department of Computational Biology and Medical Sciences, Graduate School of Frontier Sciences, The University of Tokyo, Tokyo, Japan; Division of Protein Metabolism, The Institute of Medical Science, The University of Tokyo, Tokyo, Japan; Protein Metabolism Project, Tokyo Metropolitan Institute of Medical Science, Tokyo, Japan; CIRI-International Center for Infectiology Research, Inserm, U1111, Université Claude Bernard Lyon 1, CNRS, UMR5308, Ecole Normale Supérieure de Lyon, Université Lyon, Lyon, France; Université Côte d’Azur, INSERM, C3M, 06204 Nice, France

## Abstract

RNA-protein interactions (RPIs) are as important as protein-protein interactions (PPIs) for the formation of membraneless organelles (MLOs) and play a vital role in various biological processes. Despite remarkable advances in PPI analysis technologies in recent years, the development of RPI analysis tools has lagged behind. To advance RPI analysis, we integrated three established PPI tools—bimolecular fluorescence complementation (BiFC), NanoBiT, and split-TurboID—with the RNA-targeting CRISPR/Cas13. We applied these tools to analyze paraspeckles, one of the best known MLOs formed by interactions between the long non-coding RNA NEAT1 and the RNA-binding protein NONO. The optimized BiFC-dCas13 allows live cell imaging and quantitative detection of the NEAT1-NONO interaction. The NanoBiT-dCas13 detects dynamic changes in the NEAT1-NONO interaction in an immediate and reversible manner. As a proximity labeling tool, the Split-TurboID-dCas13 induces biotinylation of proteins surrounding paraspeckles, leading to the identification of the N6-methyladenosine reader protein YTHDC1 as a novel paraspeckles-associated protein. The BiFC-dCas13, NanoBiT-dCas13, and Split-TurboID-dCas13 systems have a broad utility for the analysis of RPIs and MLOs.

## Introduction

Cells are composed of a variety of molecules, including nucleic acids and proteins. Some of these molecules assemble through liquid-liquid phase separation (LLPS) into biological condensates known as MLOs. The past decade has seen a surge in research activity surrounding LLPS and MLOs. Many MLOs are composed of RNA and RNA-binding proteins, highlighting the important role of RNA-protein interactions (RPIs) in MLO formation^1^. Therefore, the development of analytical tools to study RPIs holds immense potential for advancing our understanding of MLOs and their underlying biological functions.

Bimolecular fluorescence complementation (BiFC) is a technology that allows the detection of protein-protein interactions (PPIs) by fluorescence, in which the complementation of two split fluorescent protein fragments is induced by the interaction between proteins of interest^2^. Similarly, the luminescence reporter system based on split luciferase, called “NanoBiT”, is a robust technology for the detection of dynamic PPIs in living cells^3^. Both technologies employ signals that can be detected using tools that are commonly available in a basic laboratory setting, including fluorescence microscopy and microplate readers. Therefore, BiFC and NanoBiT are applicable to a wide range of biomedical research. Proximity labeling, such as BioID, is another invention in the field of PPI research in the last decade. Combined with immunoprecipitation followed by mass spectrometry analysis, proximity labeling allows us to investigate the PPI networks surrounding MLOs, where the molecular interactions of components are highly fluid, i.e., often fine and temporal. Split-TurboID is the pair of split fragments of biotin ligase (BL), which is advantageous for the detection of complexes induced by PPIs of two specific proteins of interest. While these technologies are highly applicable in many contexts, their use is currently limited to PPI analysis. Although RPIs are just as important as PPIs in MLOs, no analytical tools have yet been developed to investigate RPIs.

Cas13 is a CRISPR protein that targets RNA^4^ and has been widely used as a highly specific RNA knockdown tool. In addition, catalytically inactive Cas13, referred to as dCas13, has become a powerful tool to study RNA biology. In a previous study, dCas13 combined with fluorescent protein was established for tracking RNA in living cells^5^. Another study also developed dCas13 combined with BL to detect proteins surrounding specific RNA^6^. While these technologies have opened new venues for RNA research in living cells, they were not designed to detect or analyze specific RPIs. To overcome this limitation, we combined several established PPI tools with the RNA-targeting CRISPR/Cas13 and developed novel technologies: BiFC-dCas13, NanoBiT-dCas13, and Split-TurboID-dCas13. As a proof-of-concept, we applied these tools to the analysis of paraspeckle, one of the best-known MLOs in the nucleus consisting of RNA (e.g., NEAT1) and proteins (e.g., NONO). Our novel CRISPR/Cas13-based RPI analysis tools successfully detected the NEAT1-NONO interaction, visualized paraspeckles in living cells, and identified novel paraspeckle-associated proteins, including the m6A reader protein YTHDC1.

## Results

### Generation of the BiFC-dCas13 system

First we generated a BiFC-dCas13 system to visualize NEAT1-NONO interaction in paraspeckles. In this system, the split mVenus is regenerated when *NEAT1*-targeting dCas13 interacts with NONO (Figure 1A). We chose CasRX for the development of BiFC-dCas13 because CasRX has been shown to have the highest knockdown efficiency among all other Cas13s, indicating that CasRX has the strongest RNA binding ability^7^. We designed the first generation BiFC-dCas13 as a simple fusion protein, with catalytically dead CasRX (dCas13) fused directly to an N-terminal fragment of Venus split at amino acid residue 155 (VN155) at the C-terminus (Figure 1B). As a complementary partner of the Venus fragment, a C-terminal fragment of Venus split at amino acid residue 155 (VC155) was fused to NONO, also at the C-terminus. We transfected these constructs (dCas13-VN155, crRNA, and NONO-VC155) of the BiFC-dCas13 system (BiFC-dCas13_v1) into 293T cells. Live cell imaging of 293T cells transduced with the BiFC-dCas13_v1 visualized mVenus fluorescent foci with crNEAT1, which were assumed to be paraspeckles (Figure 1C). However, the mVenus signal was also detected with crNT (Figure 1C and 1D), suggesting the presence of non-specific binding of dCasRX to RNA, as shown in a previous report^5^. Since several reports have shown that the Cas13 fusion protein is most active when it includes the effector domain within its structure^8,9^, we next designed a new BiFC-dCas13 containing VN155 or VC155 within the dCas13d peptide sequence (Figure 1E). The protein structure deep learning model generated by AlphaFold^10^ showed that both dCas13 and VN155 structures are independent, even when VN155 is placed within the dCas13 peptide sequence (Figure EV1A and EV1B). In addition, the VN155 domain in the new BiFC-dCas13 is more distant from the center of the fusion protein than in the old version (Figure EV1C and Appendix Figure S1A). This indicated that the effector of the new version can operate freely from the dCas13 domain. We also replaced the SV40 NLS in the BiFC-dCas13_v1 with bipartite NLS (BPNLS), and added a double-stranded RNA-binding domain (dsRBD), which has been shown to promote the interaction between dCas13 and its target RNA^6^. Thus, we developed BiFC-dCas13_v2 as a 2nd generation BiFC-Cas13d system (Figure 1E). We transfected this BiFC-dCas13_v2 into 293T cells together with crNT or crNEAT1 and found the mVenus fluorescent foci only with crNEAT1, whereas no strong mVenus signals were seen with crNT (Figure 1F). The mean fluorescence intensity of Venus was also significantly higher in BiFC-dCas13_v2-expressing cells with crNEAT1 than those with crNT (Figure 1G). Thus, the structurally improved BiFC-dCas13, BiFC-dCas13_v2, achieved high specificity in signal-to-noise (S/N) ratio compared to the old version (Appendix Figure S1B).

**Figure 1.**
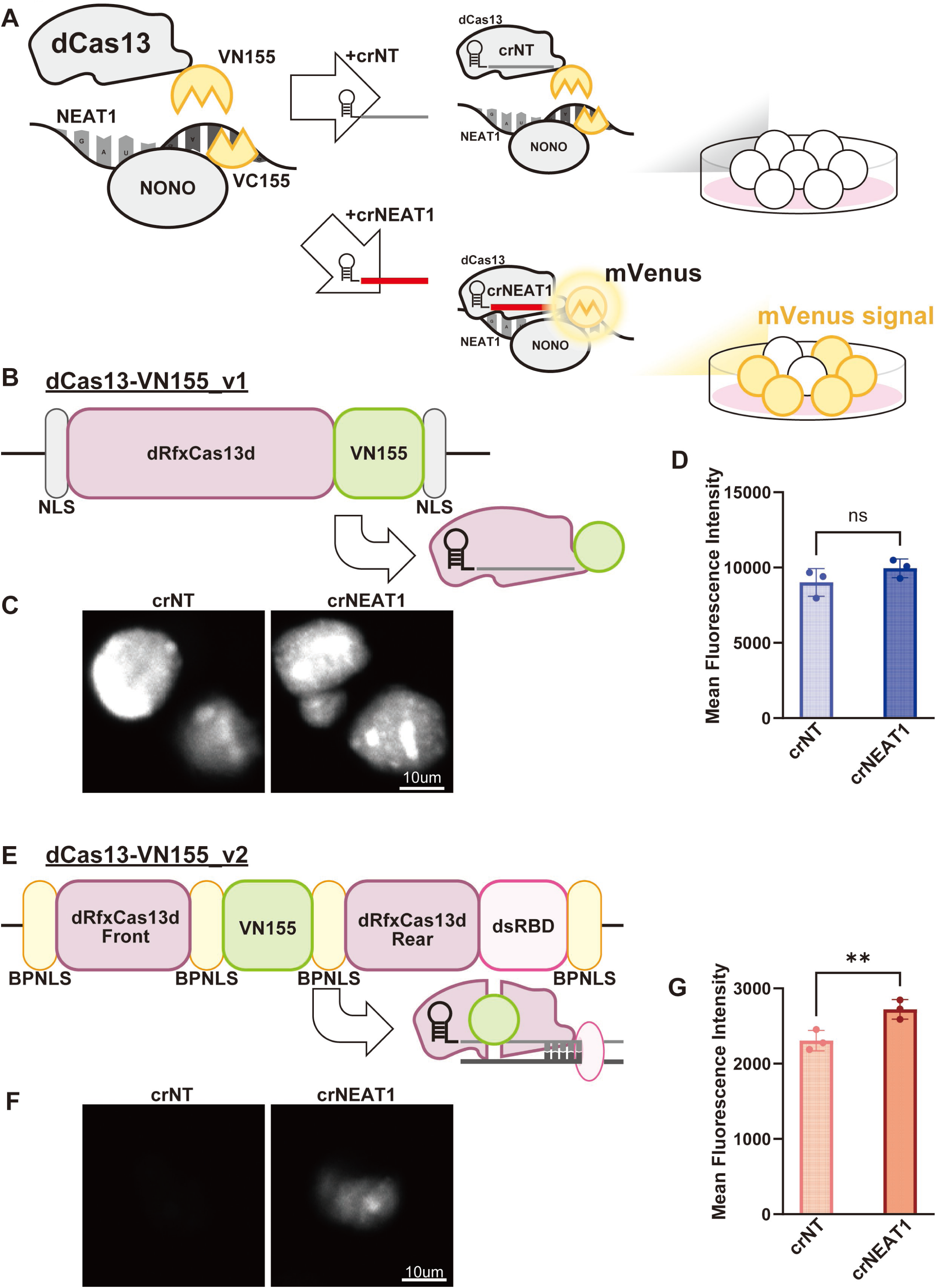
Structural improvement of the BiFC-dCas13 system. A. Schematic of BiFC-dCas13. VN155 and VC155 are fused to dCas13d and protein of interest (POI), respectively, which is NONO in this study. dCas13d-VN155 is directed to RNA of interest (ROI), which is NEAT1 in this study. BiFC complementation is induced with crRNA complementary to ROI, resulting in mVenus fluorescence glow. On the other hand, crNT, which is unable to direct dCas13d to ROI, does not induce BiFC complementation. B. Schematics of dCas13d-VN155 version 1. The VN155 domain was fused directly to dCas13d at its C-terminus. SV40 NLS was located at both ends of the fusion protein. Flexible linker (GGGGSGGGGS) was located between dCas13d and VN155. C. Live cell imaging of the BiFC-dCas13_v1 system combined with crNT or crNEAT1. 293T cells were transfected with BiFC-dCas13_v1 components and crRNAs. mVenus fluorescence images were acquired 2 days after transfection using the Nikon A1 confocal microscopic imaging system. Representative YFP fluorescence images of crNT (left), crNEAT1 (right) are shown. Scale bars: 10 μm. D. Mean fluorescence intensity of the BiFC-dCas13d_v1 systems are shown as single data points and mean +- SDs. 293T cells were transfected with BiFC-dCas13d components and crNT or crNEAT1. mVenus fluorescence signal was acquired by FACS two days after transfection. See also Appendix Figure S1B. E. Schematics of dCas13d-VN155 version 2. In version 2, dCas13d was truncated to the anterior (1-558 aa) and posterior (588-last aa) domains and VN155 was inserted between these two fragments. The double-stranded RNA binding domain (dsRBD) was fused at the C-terminus. SV40 NLS was replaced by BPNLS. Flexible linker (GGGGSGGGGS) was also placed between dCas13d and VN155 domains. F. Live cell imaging of the BiFC-dCas13_v2 system combined with crNT or crNEAT1. 293T cells were transfected with BiFC-dCas13_v2 components and crRNAs. mVenus fluorescence images were acquired 2 days after transfection using the Nikon A1 confocal microscopic imaging system. Representative YFP fluorescence images of crNT (left), crNEAT1 (right) are shown. Scale bars: 10 μm. G. Mean fluorescence intensity of the BiFC-dCas13d_v2 systems are shown as single data points and mean +- SDs. 293T cells were transfected with BiFC-dCas13d components and crNT or crNEAT1. mVenus fluorescence signal was acquired by FACS two days after transfection. See also Appendix Figure S1B.

Second, we optimized the effective NEAT1-targeting crRNAs from the entire NEAT1 sequence using the previously developed prediction model^11^ (Figure 2A) and found several effective crRNAs to visualize NEAT1-NONO interaction (Figure 2B). Cas13 can target different RNA sequences by processing tandemly arrayed premature crRNAs (spacer and 36nt direct repeat: DR36) into mature crRNAs (spacer and 30nt direct repeat: DR30)^7^. Therefore, we investigated whether arrayed crRNAs composed of multiple effective crRNAs targeting different NEAT1 sequences could improve the S/N ratio in our system (Figure 2C). As expected, arrayed crRNAs targeting multiple NEAT1 sequences showed higher signal intensity compared to single crRNAs in both flow cytometry analysis (Figure 2D) and live cell imaging (Figure 2E, 2F, and 2G). We then constructed triple crNEAT1, which showed the highest Venus reconstitution ability with the BiFC-dCas13_v2 system (Figure EV2A, 2B, and 2C). These were used in subsequent experiments.

**Figure 2.**
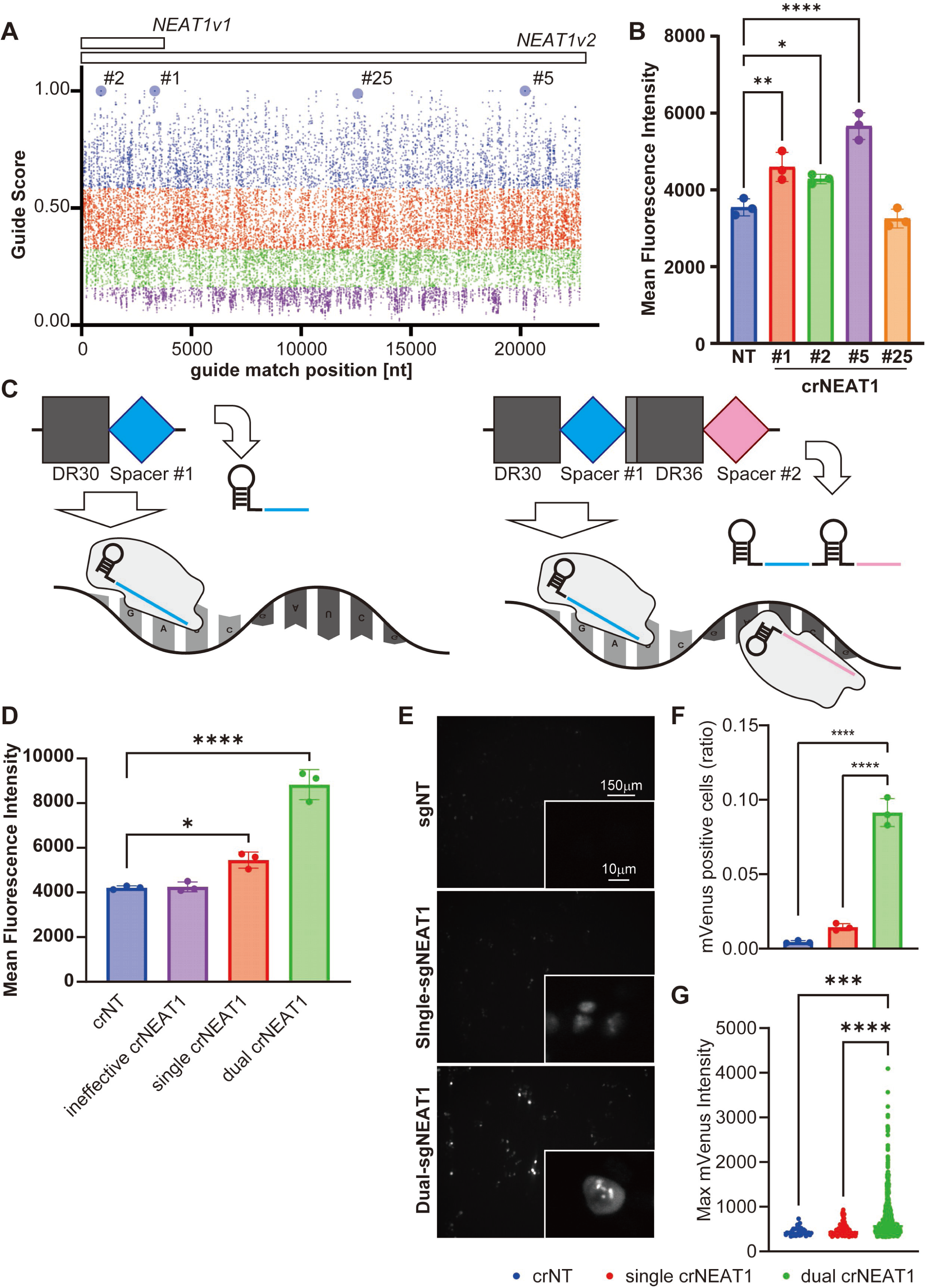
Improvement of the crRNAs used in the BiFC-dCas13 system. A. Guide scores of each crRNA targeting NEAT1 transcript position are shown. Guide scores were calculated using Cas13design (https://cas13design.nygenome.org/). Among several transcript variants, the longest isoform of NEAT1: ENST00000501122 was selected for guide score screening. 23nt crRNAs were designed from the whole transcript sequence and scored according to Cas13design criteria. Large circles indicate the crRNAs used in validation assay. B. Mean fluorescent intensity of version 2 BiFC-dCas13d systems with 5 crRNAs are shown as mean +- SDs. 293T cells were transfected with version 2 BiFC-dCas13d components and crNT, or 6 different types of crNEAT1 as indicated in Figure 2A. mVenus fluorescent signal is measured two days after transfection by FACS. *p<0.05; ****p<0.0001 (one-way ANOVA with Tukey-Kramer’s post-hoc test). N=3 for each group. C. Schematics of single and dual-crRNA, and CRISPR/Cas13 proteins guided to target transcripts. For dual-crRNA, 2 crRNAs targeting different position of same transcripts are tandemly placed. Each spacer sequences are followed by direct repeat (DR30 for former and DR36 for later crRNA) and are processed into mature crRNAs. D. Mean fluorescent intensity of version 2 BiFC-dCas13d systems with 4 crRNAs are shown as mean +- SDs. 293T cells were transfected with version 2 BiFC-dCas13d components and crNT, or three different types of crNEAT1. mVenus fluorescent signal is measured two days after transfection by FACS. *p<0.05; ****p<0.0001 (one-way ANOVA with Tukey-Kramer’s post-hoc test). N=3 for each group. E. Live cell imaging of the BiFC-dCas13d_v2 system combined with crNT, single-crNEAT1 and dual-crNEAT1. 293T cells were transfected with version 2 BiFC-dCas13d components and crRNAs. mVenus fluorescence images were acquired 2 days after transfection using the EVOS microscope imaging system. Representative Venus fluorescence images of crNT (top), single-crNEAT1 (middle), and dual-crNEAT1 (bottom) are shown (D). Scale bars: 10 μm and 150 μm. F. Ratios of mVenus-positive cells are shown as individual data points and mean +- SD. Ratios were calculated by counting the number of Venus-positive cells (above threshold) and total cells from 3 independent fields. ****P < 0.0001 (one-way ANOVA with Tukey-Kramer post hoc test). N=1. G. Maximum mVenus signal intensities of each mVenus-positive cell are shown as individual data points. Signal intensities were measured from 36, 122, and 769 cells, respectively. ***P < 0.001; ****P < 0.0001 (one-way ANOVA with Tukey-Kramer post hoc test).

Third, we constructed multiple dCas13 and NONO proteins combined with split mVenus (VN155 or VC155) at the N- or C-terminus to find the best combination with the highest S/N ratio (Appendix Figure S2A). All pairs of BiFC-dCas13 and BiFC-NONO showed complementation of the fluorescent protein when cotransfected with NEAT1-targeting crRNA in 293T cells. Among them, the combination of dCas13 with VC155 (dCas13-VC155) and NONO with VN155 at its N-terminus (VN155-NONO) showed the highest S/N ratio (Figure 3A), as indicated by the low mVenus signal in the crNT control group and the highest mVenus signal in the crNEAT1 group (Appendix Figure S2B). Live cell imaging also demonstrated that 293T cells transfected with dCas13-VC155, VN155-NONO, and crNEAT1 showed the strongest signal intensity (Figure 3B, 3C), indicating that dCas13-VC155 and VN155-NONO is the best combination. With these improvements, we have successfully established the 3rd generation BiFC-dCas13 (BiFC-dCas13_v3; Figure 3D).

**Figure 3.**
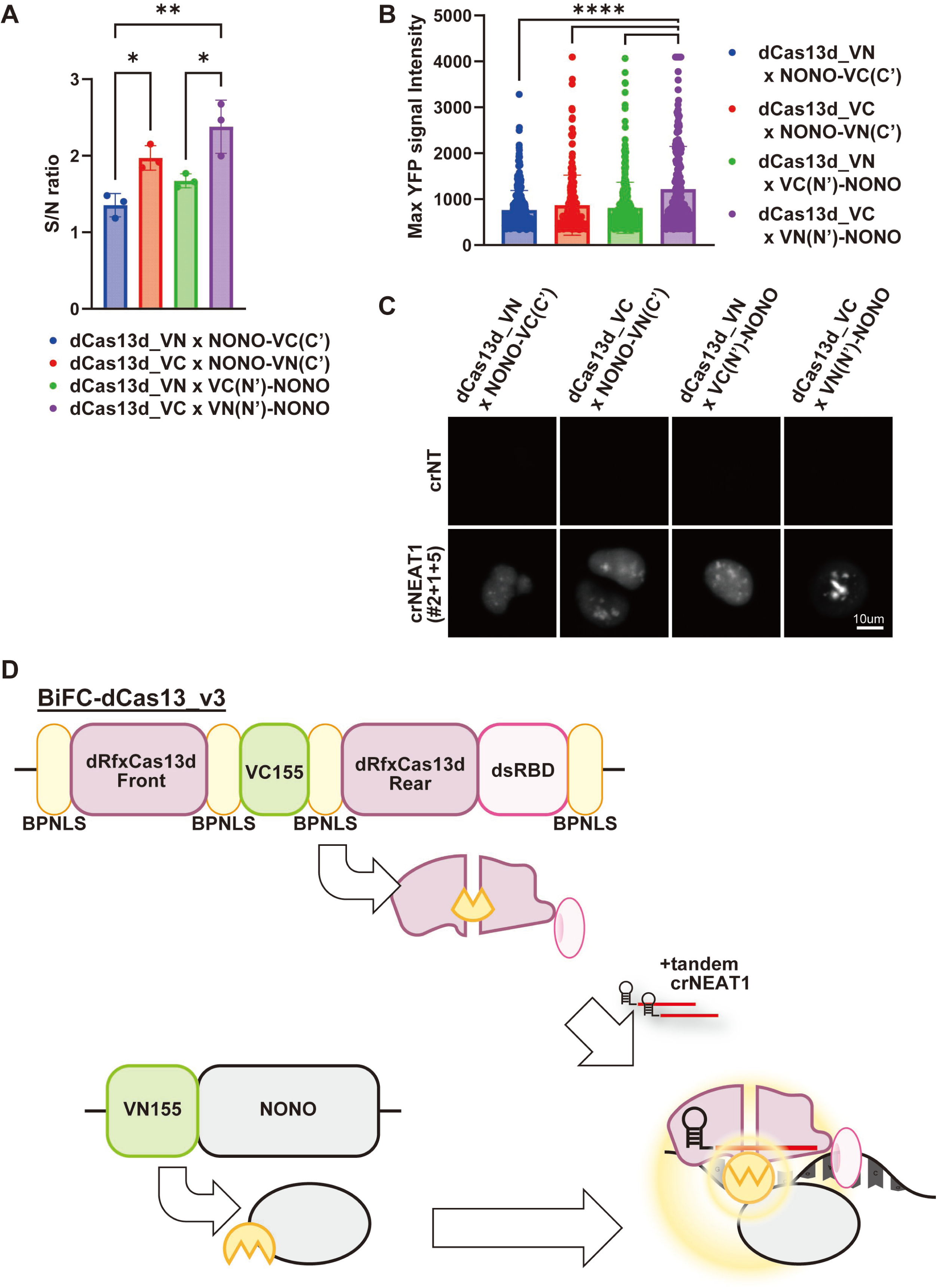
Combinational optimization between dCas13 and NONO in the BiFC-dCas13 system. A. S/N ratio of 4 different combinations of BiFC-dCas13d systems with crRNA are shown as single data points and mean +- SD. 293T cells were transfected with crRNA and BiFC-dCas13d components in different combinations as follows, dCas13d-VN155_v2 and NONO-VC155 (NONO-VC155(C’)), dCas13d-VC155_v2 and NONO-VN155 (NONO-VN155(C’)), dCas13d-VN155_v2 and VC155-NONO (VC155(N’)-NONO), and dCas13d-VC155_v2 and VN155-NONO (VN155(N’)-NONO). mVenus fluorescence signal is acquired by FACS 2 days after transfection. S/N ratio was calculated by comparing the Mean fluorescent intensity of crNEAT1 and crNT. N=3 for each group. NT, non-targeting crRNA; Du, dual-crNEAT1; Tri, triple-crNEAT1. See also Appendix Figure 2B. B. Live cell imaging analysis of the 4 different combinations of BiFC-dCas13d systems with crNT or triple-crNEAT1. 293T cells were transfected with crRNA and BiFC-dCas13d components in different combinations. mVenus fluorescence images were acquired 2 days after transfection using the Nikon-A1 confocal microscope imaging system. Maximum mVenus signal intensities are shown as individual data points and mean +-SDs. Signal intensities were measured from 293, 268, 366, and 264 cells, respectively. ****P < 0.0001 (one-way ANOVA with Tukey-Kramer post hoc test). N=1. C. Representative Venus fluorescence images of the 4 different combinations of BiFC-dCas13d systems with crNT and triple-crNEAT1 are shown. Scale bars: 10 μm. D. Schematic of the BiFC-dCas13_v3 system, which is the combination of dCas13-VC155-VN155-NONO and tandem crNEAT1.

Finally, we optimized the stoichiometry of the components of the BiFC-dCas13 system. Transfection of dCas13, NONO, and crRNA with a ratio of 1:3:2 (weight ratio) resulted in the highest mVenus expression when crNEAT1 was used, while there was no significant change in mVenus expression when crNT was used as a control (Figure EV3A).

### BiFC-dCas13 successfully detects paraspeckle formation

To confirm whether the mVenus signal of our BiFC-dCas13 system actually detects NEAT1-NONO interaction in paraspeckles, we performed NONO immunofluorescence coupled with NEAT1 FISH in 293T cells expressing the BiFC-dCas13_v3 (Figure 4A, and Appendix Figure S3A). The mVenus signals induced by crNEAT1 were correlated with NEAT1 and NONO localization (Figure 4B, and Appendix Figure S3B), and were highly enriched in NEAT1/NONO colocalized foci (Figure 4C). Thus, our BiFC-dCas13 system with crNEAT1 successfully visualized NEAT1-NONO interaction in paraspeckles.

**Figure 4.**
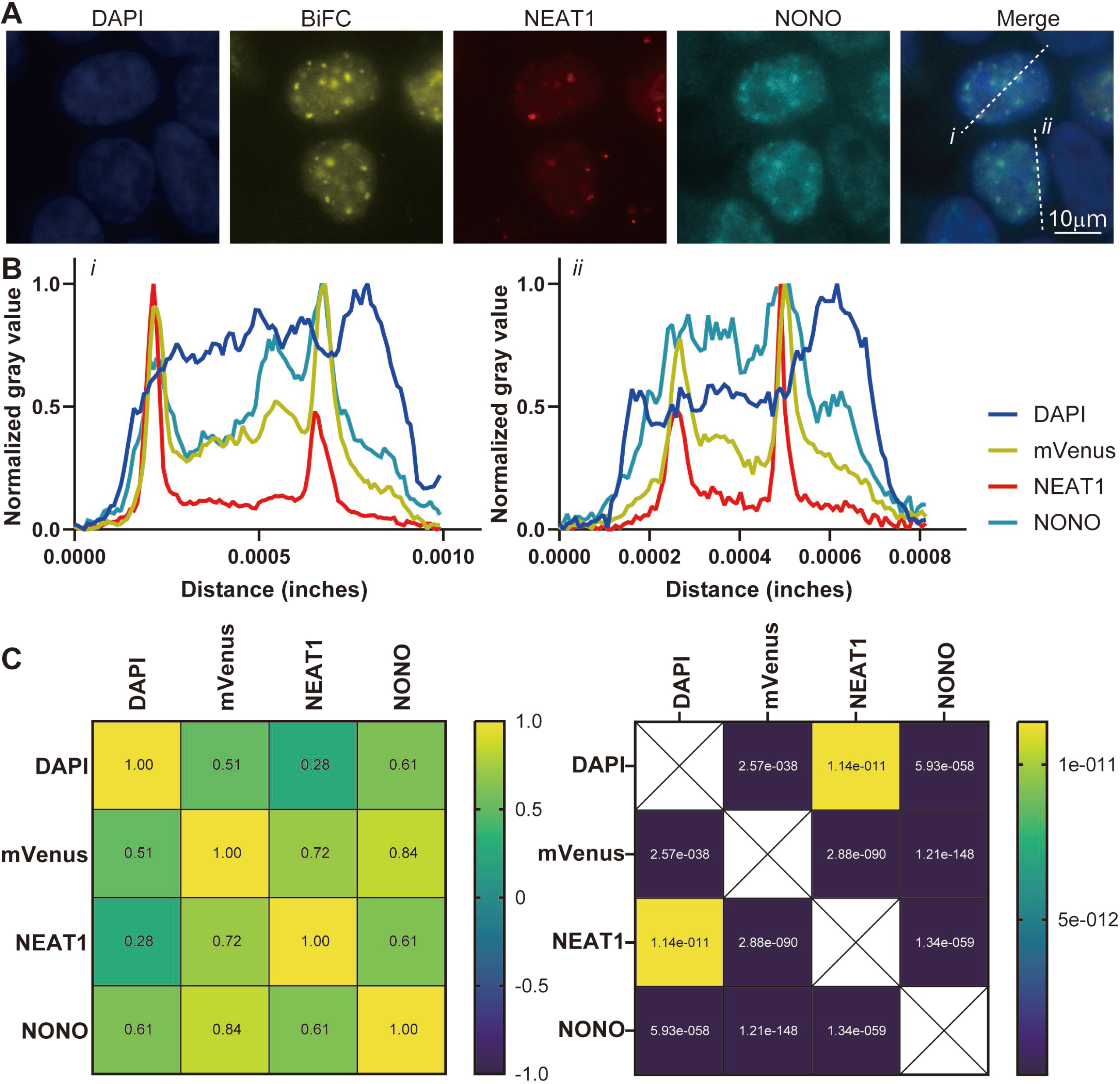
BiFC-dCas13 system visualizes paraspeckles formation. A. RNA-FISH and immunofluorescence image of 293T cells transfected with the optimized BiFC-dCas13d system. 293T cells were transfected with optimized BiFC-dCas13d components and fixed 2 days after transfection. Fluorescence images were acquired using the EVOS microscope imaging system. NEAT1_2 (red) is superimposed on mVenus (yellow), NONO (cyan) and nuclei stained with DAPI (blue). Representative images are shown. Scale bars: 10μm. See also Appendix Figure S3A. B. Plot profile analysis of signal intensity after min-max normalization. Signal intensities of DAPI (blue), mVenus (yellow), *NEAT1* (red), and NONO (cyan) were measured on the dotted lines i and ii shown in Figure 4A. See also Appendix Figure S3B. C. Colocalization analysis of DAPI, mVenus, NEAT1 and NONO. Correlations of individual signal intensities on the dotted lines, including those in Figure 4A and Figure S6A, were quantified by Pearson’s correlation. Data are presented as Pearson’s R-scores and p-values obtained from 6 cells.

### Generation of the NanoBiT-dCas13 system

One limitation of the BiFC-dCas13 system is its irreversibility. Because the reconstituted fluorochrome is essentially irreversible, the system cannot monitor the dynamic change of RPI^12^. To overcome this limitation, we developed an alternative reporter system using NanoBiT technology. NanoBiT is a split luciferase consisting of LgBiT and SmBiT that functions as NanoLuc only when these two molecules are in proximity to each other^3^. Importantly, the interaction between LgBiT and SmBiT is reversible, allowing this technology to be used to monitor dissociating proteins.

We developed the NanoBiT-dCas13 system, in which the N- and C-terminal mVenus fragments were replaced by the LgBiT and SmBiT. In this system, the LgBiT and SmBiT interaction is induced by dCas13 targeting a specific RNA and protein of interest (Figure 5A). Two combinations of NanoBiT-dCas13, LgBiT-dCas13, and NONO-SmBiT, or vice versa, were generated and transduced together with crRNAs in 293T cells. Combined expression of dCas13-LgBiT and NONO-SmBiT in conjunction with crNEAT1 resulted in the production of robust luminescence signals, whereas the combination of dCas13-SmBiT and NONO-LgBiT did not (Figure 5B). Similar to BiFC-dCas13, the luciferase activity of NanoBiT-dCas13 with multiple crNEAT1 was higher than that with single crNEAT1 (Figure 5B). We investigated the reversibility of the NanoBiT-dCas13 system using 1,6-hexandiol (1,6-HD), which is a widely used reagent to dissolve LLPS-forming MLOs^13^. Five minutes incubation with 5% 1,6-HD completely abolished the luciferase activity of NanoBoT-dCas13 with crNEAT1 (Figure 5C), demonstrating the reversibility of this system. To investigate the stimuli that affects paraspeckle formation, we exposed 293T cells transfected with NanoBiT-dCas13 to various reagents (Appendix Figure S4). Interestingly, a BCL2 inhibitor venetocrax^14^ enhanced the NEAT1-NONO interaction (Figure 5D), suggesting the possible involvement of BCL2 in paraspeckle formation. Thus, we successfully developed the NanoBiT-dCas13 system to detect paraspeckle formation in a reversible manner.

**Figure 5.**
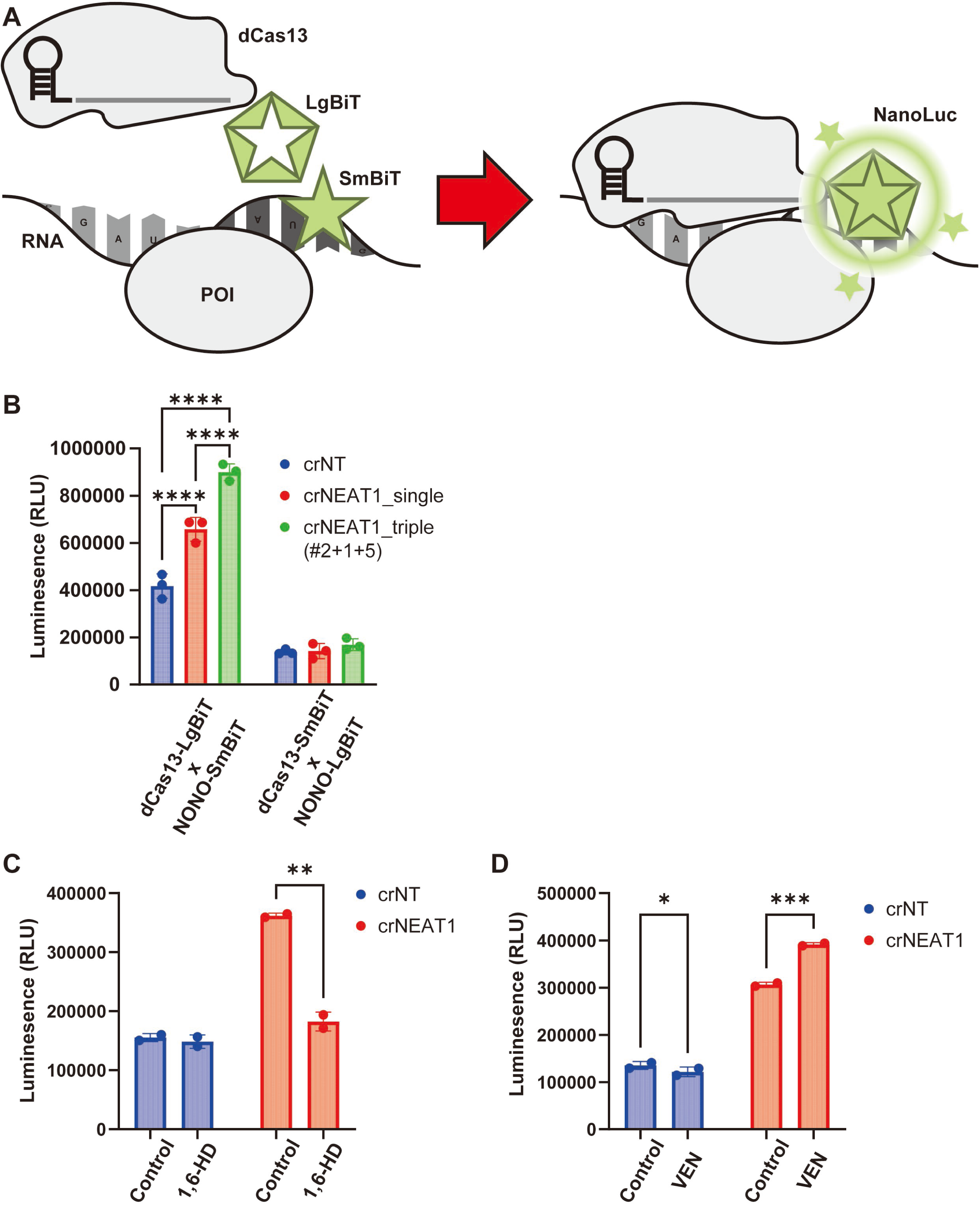
NanoBiT-dCas13 system detects paraspeckle formation in a reversible manner. A. Schematic of NanoBiT-dCas13d. LgBiT and SmBiT are fused to dCas13d and protein of interest (POI), respectively. dCas13d-LgBiT is directed to RNA of interest (ROI) with crRNA and luciferase complementation is induced. When the complemented luciferase acts on the luciferin substrate, luminescence is emitted. B. Luminescence intensity of different combinations of NanoBiT-dCas13d systems with crRNAs are shown as individual data points and mean +- SD. 293T cells were transfected with NanoBiT-dCas13d components and crRNAs. Nano-Glo luciferin substrate was added, and luminescence was measured 2 days after transfection using a microplate reader. N=3 for each group. ****P < 0.0001 (one-way ANOVA with Tukey-Kramer post hoc test). C. Luminescence intensity of NanoBiT-dCas13d systems with crRNAs under the condition of 1,6-HD are shown as individual data points and mean +- SD. 293T cells were transfected with NanoBiT-dCas13d components and crRNAs. 2days after the transfection, cells were treated with 5% 1,6-HD for 5 minutes and washed out. Nano-Glo luciferin substrate was added, and luminescence was measured using a microplate reader. N=2 for each group. **p<0.01 (unpaired t test) D. Luminescence intensity of NanoBiT-dCas13d systems with crRNAs under the condition of VEN are shown as individual data points and mean +- SD. 293T cells were transfected with NanoBiT-dCas13d components and crRNAs. 2days after the transfection, cells were treated with 10mM of VEN for 30 minutes and washed out. Nano-Glo luciferin substrate was added, and luminescence was measured using a microplate reader. N=2 for each group. *p<0.05; ***p<0.001 (unpaired t test)

### Generation of the Split-TurboID-dCas13 system

Finally, we developed the Split-TurboID-dCas13 system to study the interactomes surrounding specific RNA-protein complexes. In this system, N- and C-terminal fragments of TurboID, TbN73 and TbC74, are reconstituted in response to specific RPI and biotinylate surrounding proteins in the presence of biotin (Figure 6A). We constructed fusion proteins of dCas13 and NONO with N- or C-terminal TurboID, generating several dCas13-split TurboID fusions (dCas13-TbN73_v2 or dCas13-TbC74_v2) and NONO-split TurboID fusions (NONO-TbN73/TbC74; TbN73/TbC74-NONO). Co-transfection of dCas13-TbN73_v2, NONO-TbC74 and crNEAT1 into 293T cells induced protein biotinylation, and the biotinylated proteins were enriched by streptavidin (SA) bead immunoprecipitation (Figure 6B). We then evaluated the biotinylation-inducing activity of four sets of Split-TurboID-dCas13 system coexpressed with single, double (crNEAT1#2+1) or triple (crNEAT1#2+1+5) crNEAT1, and found that dCas13-TbN73_v2 and NONO-TbC74 together with triple crNEAT1 was the best combination to efficiently induce biotinylation of nearby proteins (Figure 6C, and Appendix Figure S5).

**Figure 6.**
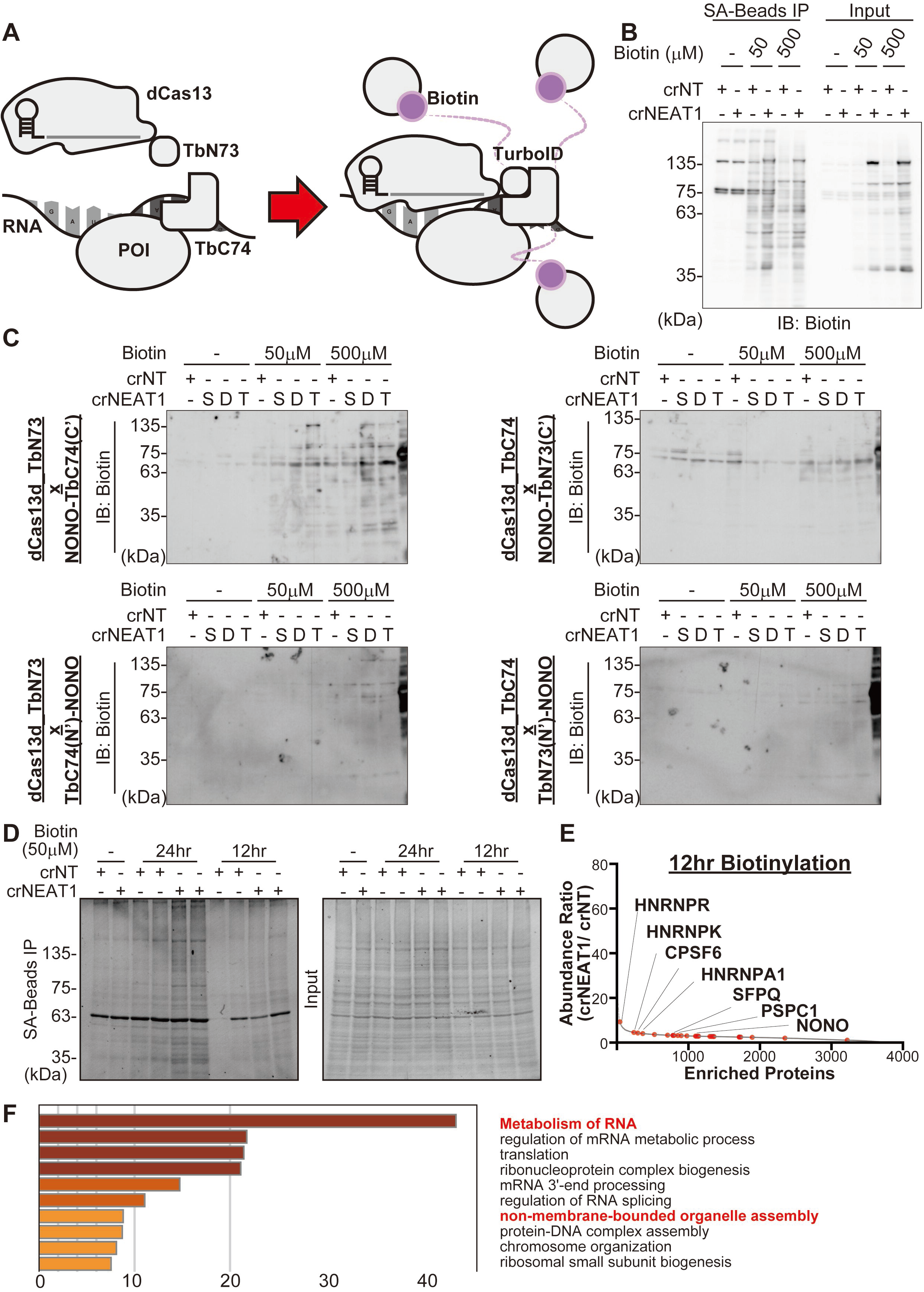
Split-TurboID dCas13 system enables proximity labeling of proteins surrounding paraspeckles. A. Schematic of NanoBiT-dCas13d. TbN73 and TbC74 are fused to dCas13d and protein of interest (POI), respectively. dCas13d-TbN73 is directed to RNA of interest (ROI) with crRNA and biotin ligase (BL) complementation is induced. In the presence of biotin, BL biotinylates proteins surrounding the POI-ROI complex. B. Representative biotin immunoblot of streptavidin immunoprecipitation from 293T transfected with the split-turboID-dCas13d system. 293T cells were transfected with dCas13d-TbN73_v2, NONO-TbC74, and crRNAs. Two days after transfection, biotinylation was induced at the indicated concentration for 24hours. Total cell lysates were isolated and immunoprecipitated with streptavidin beads, and biotinylated proteins were detected with anti-biotin antibody. C. Representative biotin immunoblot of 4 different combinations of split-TurboID-dCas13d systems. 293T cells were transfected with crRNA and BiFC-dCas13d components in different combinations as follows, dCas13d-TbN73_v2 and NONO-TbC74 (NONO-TbC), dCas13d-TbC74_v2 and NONO-TbN73 (NONO-TbN), dCas13d-TbN73_v2 and TbC74-NONO (TbC-NONO), and dCas13d-TbC74_v2 and TbN73-NONO (TbN-NONO). Two days after transfection, biotinylation was induced at the indicated concentration for 24hours. Total cell lysates were subjected to Western blot with anti-biotin antibody. S, single-crNEAT1; D, dual-crNEAT1; T, triple-crNEAT1. D. Oriole staining after streptavidin immunoprecipitation of 293T cells transfected with optimized split-TurboID-dCas13d systems. 293T cells were transfected with dCas13d-TbN73_v2, NONO-TbC74 and crNT or triple-crNEAT1. Two days after transfection, cells were incubated with 50μM biotin for the indicated time course. Total cell lysates were isolated and immunoprecipitated with streptavidin beads. Enriched proteins were analyzed by oriole staining. E. Fold change (crNEAT1/ crNT) in protein abundance. Immunoprecipitated proteins after 12 hours of biotinylation induced by split-TurboID-dCas13d systems were subjected to LCMS/MS analysis. Red dots indicate proteins known as paraspeckle components^15^. N=2 for each group. See also Figure EV4B. F. Enriched ontology clusters of the split-TurboID-dCas13d with crNEAT1. Enriched proteins in the crNEAT1 group (FDR<0.2 in both biotinylations) were analyzed by Metascape^16^.

Using this Split-TurboID-dCas13 system, we next performed an interactome analysis around the NEAT1-NONO complex in paraspeckles. We transfected constructs of the Split-TurboID-dCas13 system into 293T cells and induced biotinylation for 12 or 24 hours. Biotinylated proteins were then collected using SA beads and subjected to liquid chromatography-tandem mass spectrometry (LC-MS/MS) (Figure 6D). Similar proteins were biotinylated at both 12 and 24 h (Figure EV4A). LC-MS/MS analysis revealed that known paraspeckle proteins, such as SFPQ and PSPC1^15^, were biotinylated in crNEAT1-transduced 293T cells (Figure 6E, and Figure EV4B). Enrichment analysis using Metascape^16^ showed that proteins related to RNA metabolism and those that assemble non-membrane-bound organelles were highly enriched (Figure 6F). These results suggest that paraspeckle is closely associated with RNA regulation, as has been shown in previous reports^17^.

### YTHDC1, an m6A reader protein, supports paraspeckle formation

Interestingly, the Split-TurboID-dCas13 system with crNEAT1 showed that m6A RNA regulatory proteins, especially m6A “reader” proteins, were highly enriched in biotinylated proteins (Figure EV5A). Among these, we focused on YTHDC1 because it localizes to the nucleus where paraspeckle forms. To investigate the possible interaction between paraspeckle and YTHDC1, we established a reporter cell line to visualize endogenous YTHDC1 and a paraspeckle protein NONO with mNeonGreen and mCherry, respectively (Figure EV5B). We first introduced the sequence of the small mNeonGreen fragment (mNG_2_11)^18^ into the C-terminal locus of the *NONO* gene to express the NONO-mNG_2_11 fusion protein in K562 cells expressing the large mNeonGreen fragment [mNG_2_(1-10)]. In this reporter cell line, the NONO-mNG_2_11 complements with mNG_2_(1-10) to restore fluorescence reflecting NONO expression. We also introduced the mCherry sequence into the N-terminus locus of the *YTHDC1* gene in K562 cells and established an mCherry-YTHDC1 reporter cell line and a NONO/YTHDC1 dual reporter cell line (Figure EV5C, and EV5D). Successful knockin in these reporter cell lines was confirmed by PCR (Appendix Figure S6). Live cell imaging of the dual reporter cells revealed the broad coexistence of NONO and YTHDC1 (Figure EV5D, EV5E), indicating the close association between YTHDC1 and paraspeckles. However, compared to the signal of NONO, which forms paraspeckle foci in the nucleus, the YTHDC1 signal was flatter, more diffuse and localized “around” the peak of the NONO signal (Figure EV5E). Therefore, YTHDC1 is unlikely to be a core component of paraspeckles, but rather a co-operating protein surrounding paraspeckles.

Since m6A reader proteins regulate the fate of RNAs, we speculated that m6A readers are associated with paraspeckle formation through *NEAT1* regulation. To test this possibility, we depleted YTHDC1 and its family gene, YTHDC2, using the CRISPR/Cas9 system in K562 (Figure 7A). Depletion of YTHDC1, but not YTHDC2, induced a significant reduction of *NEAT1* expression (Figure 7B). NONO immunofluorescence coupled with *NEAT1* FISH also revealed the reduction of paraspeckles in YTHDC1-depleted K562 cells (Figure 7C, 7D). Taken together, these results suggest that YTHDC1 coexists with NONO, although not necessarily in the paraspeckle foci, and functionally supports paraspeckle formation by stabilizing NEAT1.

**Figure 7.**
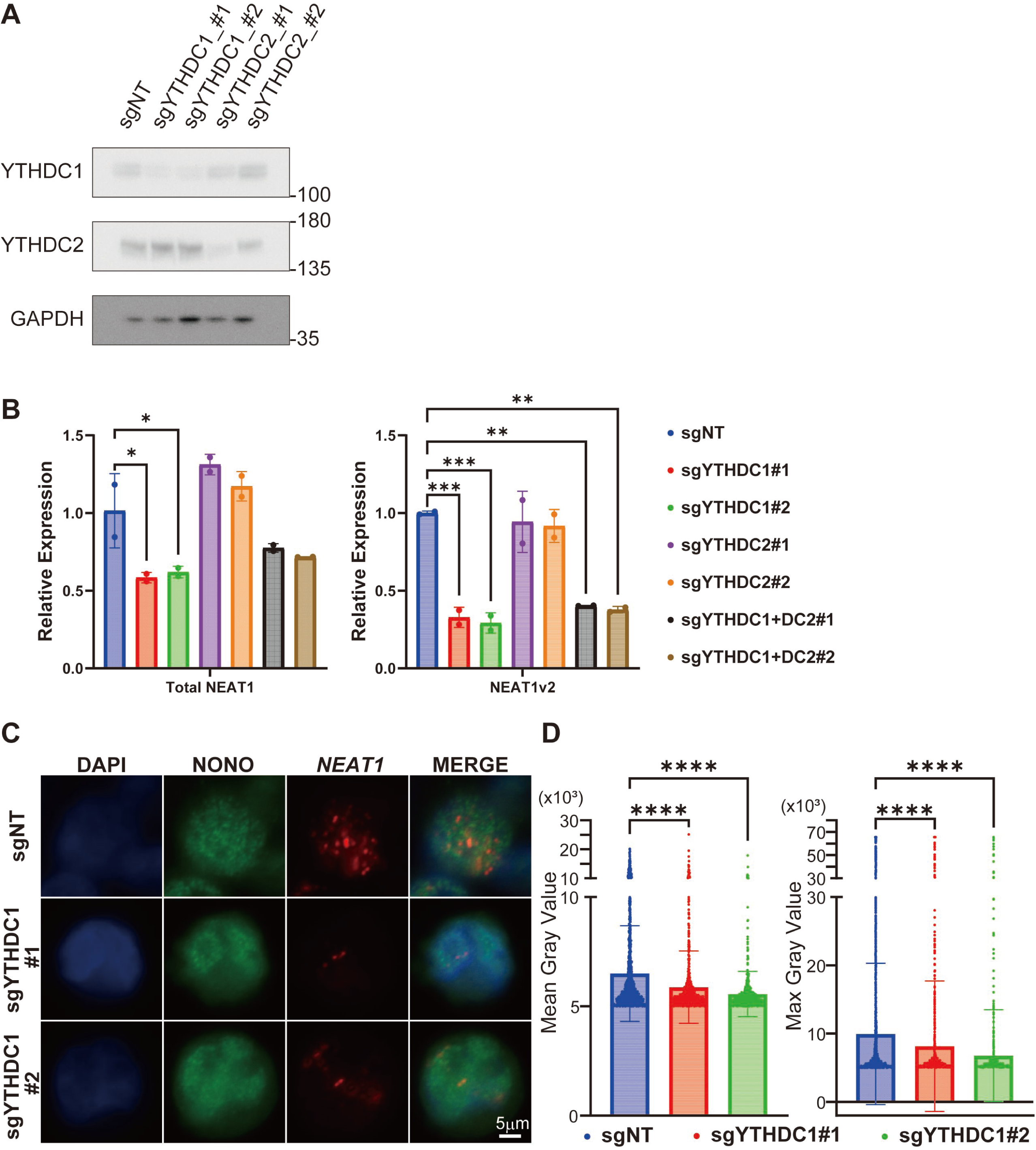
YTHDC1 supports paraspeckle formation through NEAT1 stabilization. A. Depletion of YTHDC1 and YTHDC2 in K562 cells was confirmed by Western blotting. Cells were transduced with non-targeting (NT), YTHDC1-targeting (sgYTHDC1), or YTHDC2 sgRNA (sgYTHDC2) (coexpressing puromycin resistance gene). Total protein was extracted from cells after 3 days of puromycin selection. B. Relative expression of total NEAT1 and *NEAT1* variant 2 (*NEAT1*v2) in YTHDC1 or YTHDC2 depleted K562 cells shown as means ± SDs. K562 expressing Cas9 were transduced with non-targeting YTHDC1 or YTHDC2 targeting sgRNA (co-expressing puromycin resistance gene). RNA was extracted from cells after puromycin selection and subjected to RT-qPCR. Data were analyzed after ACTB normalization. *p < 0.05; **p < 0.01; ***p < 0.001 (one-way ANOVA with Dunnett post hoc test). N = 2 (technical replicate) for each group. C. RNA-FISH and immunofluorescence image of YTHDC1 or YTHDC2 depleted K562 cells. K562 expressing Cas9 were transduced with non-targeting YTHDC1 targeting sgRNA (co-expressing puromycin resistance gene). Cells were fixed 2 days after puromycin selection. Fluorescence images were acquired using the Nikon A-1 microscope imaging system. *NEAT1*_2 (red) is superimposed on NONO (green) and nuclei stained with DAPI (blue). Representative images are shown. Scale bars: 5 μm. D. Mean (left) and maximum (right) *NEAT1* signal intensities in each cell are shown as individual data points and mean +-SDs. Signal intensities were measured from 2773, 1082, and 779 cells, respectively. ****P < 0.0001 (one-way ANOVA with Dunnett post hoc test).

## Discussion

In this work, we developed three CRISPR/Cas13-based tools to detect or analyze RPIs. As an imaging tool for RPIs, BiFC-dCas13 successfully visualize paraspeckle formation by detecting *NEAT1*-NONO interactions. BiFC-dCas13 has advantages over conventional dCas13-based imaging tools because fluorescence excitation is directly controlled by the RPI, thus overcoming the drawback of the dCas13 fluorescent protein, which can fluoresce independently of its target^5^. This property of BiFC-dCas13 allows quantitative detection of the RPIs by the fluorescence signal strength, opening a new venue for the analysis of RPIs. Another technology used for RNA tracking is MS2-MCP, in which RNAs with MS2 sequences interact with MCP^19^. Because MS2 sequences must be present in the target transcript, this system is not suitable for tracking endogenous RNA. Combined with the ease of crRNA design, our BiFC-dCas13 system provides an easy and rapid approach to visualize endogenous RPIs in living cells.

Unfortunately, the BiFC-dCas13 system is unlikely to be useful for drug screening approaches due to its irreversibility. We have therefore developed the NanoBiT-dCas13 system, which can quantify the strength of RPI using a microplate reader. Because NanoBiT-dCas13 is a reversible system, it will be highly affinitive for high-throughput screening (HTS) using CRISPR-based gRNA libraries and/or compound libraries, which may lead to a paradigm shift in the investigation of new drugs targeting endogenous RPIs in the future.

MLOs are composed of numerous molecules and play important roles in diverse biological processes. Our Split-TurboID-dCas13 system is highly effective in identifying novel interacting proteins in various types of MLOs. We applied this tool to investigate the interactome of paraspeckle and successfully identified YTHDC1 as a novel paraspeckle-associated protein. YTHDC1 is an m6A RNA reader protein and has multiple functions in cells, including RNA transfer, processing, and stabilization. In addition, a recent study showed that YTHDC1 itself forms condensate to suppress myeloid differentiation^20^. In this study, we found that YTHDC1 is present in the periphery of NONO-NEAT1 speckles to regulate NEAT1 stability, revealing the novel role of YTHDC1 in paraspeckle formation. Since proximity labelling technologies are now used to capture nucleic acids^21^, the Split-TurboID-dCas13 system can also be applied to identify DNAs and RNAs that interact with specific MLOs. Indeed, paraspeckles and NEAT1 have been shown to interact with multiple RNAs and genomic loci^22^ ^23^, which can be systematically identified using the Split-TurboID-dCas13 system.

The Cas13-based RPI analytical tools developed in this study contain multiple NLS sequences and mainly target nuclear RNAs and proteins. However, there are numerous cytoplasmic MLOs, including stress granules and P-bodies whose functions are not fully understood^24^. The development of similar tools targeting cytoplasmic MLOs is an important future challenge. To study endogenous RPIs in physiological states, it will also be important to combine the knock-in technology with our systems. For example, we can introduce split mVenus or split TurboID fragment sequences into the N- or C-terminus of the protein of interest (e.g., NONO). With this method, we will be able to efficiently detect or analyze various endogenous RPIs in living cells in future studies.

In summary, we have combined RNA-targeting dCas13 and split protein systems and developed novel analytical tools to detect or analyze RNA-protein interactions. Using these tools, we visualized NEAT1-NONO interactions in paraspeckle and identified YTHDC1 as a novel paraspeckle-associated protein. These tools and findings pave the way for future innovations in RPI studies.

## Material and methods

### Cell culture

293T cells were cultured in DMEM containing 10% fetal bovine serum (FBS) and 1% Penicillin-Streptomycin (nacalai). K562 cells were cultured in RPMI-1640 medium supplemented with 10% FBS and 1% Penicillin-Streptomycin (nacalai). Cells were cultured in humidified incubators at 37°C in an atmosphere of 5% CO_2_.

### Plasmids

All plasmids used in this study were cloned by PCR amplification and HiFi DNA assembly methods. For the construction of dCas13 fusion genes, dCas13 and dCas13-dsRBD were used as backbone vectors^6,7^. For the construction of NONO fusion genes, pMYs-IN NONO was used as the backbone vector^25^. DNA sequences of VN155/ VC155^2^, LgBiT/ SmBiT (promega)^3^, and TbN73/ TbC74^26^ were PCR amplified using Phusion High-Fidelity DNA Polymerase (New England Biolabs) or KOD FX Neo (TOYOBO) and assembled into an enzymatically restricted backbone vector using the NEB HiFi DNA Assembly Kit (New England Biolabs). For CRISPR/Cas9 knock-in, 400 bp of left and right homology arm (HA) of YTHDC1, flanked by guide RNA targeting sequence and PAM, was designed and synthesized using GeneArt Gene Synthesis (Thermo Fisher Scientific). mCherry sequence was assembled between the two HAs by PCR amplification and HiFi DNA assembly methods. NEB 5-alpha competent E. coli (New England Biolabs) was used for plasmid propagation. For crRNA production or guide RNA construction, annealed oligos were inserted into crRNA backbone vector^7^ or lentiGuide-Puro vector^27^. crRNA sequences were designed with Cas13design (https://cas13design.nygenome.org/). sgRNA sequences were designed with CRISPick (https://portals.broadinstitute.org/gppx/crispick/public). The sequences of the crRNAs and sgRNAs used in this study are described in the Appendix Table S1.

pXR002: EF1a-dCasRx-2A-EGFP (Addgene plasmid # 109050; http://n2t.net/addgene:109050; RRID:Addgene_109050) and pXR003: CasRx gRNA cloning backbone (Addgene plasmid # 109053; http://n2t.net/addgene:109053; RRID:Addgene_109053) were gifts from Patrick Hsu. 3xHA-TurboID-NLS_pCDNA3 was a gift from Alice Ting (Addgene plasmid # 107171; http://n2t.net/addgene:107171; RRID:Addgene_107171). lentiCas9-Blast (Addgene plasmid # 52962; http://n2t.net/addgene:52962; RRID:Addgene_52962) and lentiGuide-Puro (Addgene plasmid # 52963; http://n2t.net/addgene:52963; RRID:Addgene_52963) were gifts from Feng Zhang. Lentiviral packaging vectors [(pMD2.G # 12259; http://n2t.net/addgene:12259; RRID:Addgene_12259) and (psPAX2 # 12260; http://n2t.net/addgene:12260; RRID:Addgene_12260)] were gifts from Didier Trono. pDGM6 was a gift from David Russell (Addgene plasmid # 110660; http://n2t.net/addgene:110660; RRID:Addgene_110660). BaEVRless^28^ was gift from Els Verhoeyen.

### Transient transfection and virus (or virus-like particle) production

For transient transfection of BiFC-dCas13, NanoBiT-dCas13 and split-TurboID-dCas13, 293T cells at ∼50% confluence were transfected with DNA plasmid using PEI or Lipofectamine 3000 (Thermo Fisher Scientific), and subjected to further assay 1-2 days after transfection. The mixing ratio of the transfected DNA is dCas13: NONO: crRNA = 1:3:2, unless otherwise noted.

For lentivirus, adeno-associated virus (AAV) and MMLV virus-like particle (VLP)^29^, 293T cells at ∼80% confluence were transfected with DNA plasmid using the calcium-phosphate method. Lentiviruses were produced by transient transfection of 293T cells with viral plasmids together with lentiviral packaging plasmids (pMD2.G and psPAX2). AAV were produced by transient transfection of 293T cells with viral plasmids together with AAV serotype 6 packaging plasmids (pDGM6). Cas9-MMLV VLPs were produced by transient transfection of 293T cells with MMLV gag-Cas9 plasmids and sgRNA plasmids together with MMLV gag-pol and envelope expressing plasmids (VSVg and BaEVRless^30,31^). All virus (VLP) particles were harvested from the supernatant 48-72 hours post-transfection and filtered through a 0.45 um PVDF membrane filter. For Cas9-MMLV VLP concentration, 1/3 volume of Lenti-X Concentrator (Takara) was added to the supernatant, incubated overnight at 4°C, and pelleted by centrifugation at 1,500g for 45 min at 4 °C. The VLP pellet was resuspended in Opti-MEM to achieve 10-fold concentration. Viruses (or VLPs) were stored at -80°C and thawed immediately before use.

### Gene knockout using CRISPR/Cas9

For *YTHDC1* and *YTHDC2* depletion, K562 cells were transduced with Cas9 (coexpressing blasticydine deaminase) using lentiviruses produced as described above. After 1 week of blasticidin selection, K562/Cas9 cells were transduced with guide RNAs (coexpressing puromycin resistance gene). After 3 days of puromycin selection, the cells were subjected to the RNA or protein extraction or RNA-FISH analysis. The sequences of the sgRNAs used in this study are described in the Appendix Table S1.

### Gene knock-in using CRISPR/Cas9

For gene editing (knock-in) of K562 genome, donor DNA of mNG211 was prepared as ssODN (IDT). Donor DNA of mCherry was prepared as AAV-producing plasmid as described above. Donor DNA was transfected into K562 cells using Lipofectamine 3000 (ThermoFisher) or rAAV6 viral transduction, followed by target-site specific Cas9 VLP to induce homologous directed repair.

The sequences of the sgRNAs used in this study are described in the Appendix Table S1.

### Live cell imaging and RNA-FISH followed by immunofluorescence

Fluorescence images were captured using an EVOS imaging system (Invitrogen), a confocal microscope systems (A1Rsi [Nikon] or IX73 [Olympus] equipped with a spinning disk confocal unit CSU-W1 [Yokogawa]). Images were analyzed using NIS-Elements software or ImageJ Fiji image analysis software. For sample preparation, 293T cells cultured on round glass slides or K562 cells cultured on poly-L-lysin-coated glass slides were used. RNA-FISH was performed according to the manufacturer’s instructions. Briefly, cells were fixed with 4% paraformaldehyde and permeabilized with 70% ethanol. The cells were then hybridized with stellaris FISH probes, human *NEAT1* middle segment with Quasar 570 dye, and stained with anti-NONO, antibodies, followed by labeling with AlexaFluor488-conjugated anti-mouse (Invitrogen, A-11029) or AlexaFluor633-conjugated anti-rabbit (Invitrogen, A-21086) antibodies. Cell nuclei were visualized with DAPI.

### Flow cytometry

For quantification of BiFC-dCas13, the signal strength of mVenus in cells was analyzed using a CytoFlex LX (Beckman Coulter, Inc.) with CytExpert software. To isolate genome-edited K562 cells, we sorted mNeonGreen-positive, mCherry-positive, and mNeonGreen/mCherry double-positive cells using a FACSAria (BD bioscience) with FACSDiva software.

For 293T sample preparation, after transient transfection of the BiFC-dCas13 system consisting of dCas13-VN155/ VC155, NONO-VC155/ VN155 and crRNA, cells were trypsinized to detach from the dish, centrifuged and resuspended in PBS containing 2% FBS. For K562 sample preparation, gene-edited cells were harvested, centrifuged once, and resuspended in PBS containing 2% FBS. 4’, 6-diamidino-2-phenylindole (DAPI) was used to exclude dead cells.

All flow cytometry analyses were performed using Flowjo software (BD bioscience), with gating strategies shown in Appendix Figure S7.

### Luciferase assay

NanoBiT PPI luminescence was measured using the Nano-Glo Live Cell Assay System (Promega) according to the manufacturer’s instructions. Briefly, 293T cells were seeded in 96-well culture plates at a density of 10,000 cells per well. At 18 hours after seeding, the cells were transfected with the NanoBiT-dCas13 system using polyethyleneimine (PEI). Nano-Glo Live Cell Reagent was added 2 days after transfection and assayed for luciferase activity using the luciferase assay system (Promega) and a luminometer (BMG LABTECH, FLUOstar OPTIMA).

### Quantitative RT-PCR

RNA was extracted using TRIzol and reverse transcribed using High Capacity cDNA Reverse Transcription Kits (Applied Biosystems, 4368814) according to the manufacturer’s protocol. cDNA was then subjected to quantitative RT-PCR using a SYBR Select Master Mix (Applied Biosystems) with a StepOnePlus System (Applied Biosystems). The sequences of the primers used for qRT-PCR in this study are described in the Appendix Table S1.

### Biotin immunoprecipitation (IP) assay

293T cells were transiently transfected with Split-TurboID-dCas13 system using PEI. Biotinylation was induced 48 hours after transfection for the indicated time, and cells were lysed in total cell lysis buffer [50 mM Tris-HCl pH7.5, 1 mM EDTA, 150 mM NaCl, 1% Triton X-100, 2 mM PMSF, and 1 mM DTT protease inhibitor cocktail (Sigma)]. Sample lysates were incubated with Dynabeads M-280 Streptavidin (Invitrogen) for 30 minutes at room temperature. Precipitates were washed three times with cell lysis buffer and eluted in Laemmli sample buffer (Bio-Rad). Samples were subjected to sodium dodecyl sulfate-polyacrylamide gel electrophoresis (SDS-PAGE) followed by gel staining or Western blot. For Oriole staining, the gel was stained with Oriole fluorescent gel stain (Bio-Rad) for more than 2 hours at room temperature and visualized. For Western blotting, transferred proteins were stained with anti-biotin antibody (Santa Cruz Biotechnology). Signals were detected with ECL Western Blotting Substrate (Promega), and immunoreactive bands were visualized with a LAS-4000 Luminescent Image Analyzer (FUJIFILM). Band intensities were analyzed using LabWorks version 4.5 software (UVP, LLC).

### Mass-spectrometric analyses

The LC-MS/MS analysis was performed essentially as described previously with minor modifications^32^. Analytes were separated by SDS-PAGE with a short run (1 cm) and stained with Bio-Safe Coomassie (Bio-Rad). The gel pieces were washed with 50 mM ammonium bicarbonate (AMBC)/30% acetonitrile (ACN) for 30 min and with 50 mM AMBC/50% ACN for 30 min. Proteins were reduced in 10 mM dithiothreitol (DTT)/50 mM AMBC for 15 min at 56°C and alkylated in 55 mM iodoacetamide (IAA)/50 mM AMBC for 45 min at RT in the shade. The gel pieces were dehydrated in 50% ACN/50 mM AMBC for 10 min and then in 100% ACN for 5 min, followed by in-gel tryptic digestion with 20 ng/μL Trypsin Gold (Promega) in 50 mM AMBC/5% ACN at 37°C for 16 h. The digested peptides were extracted with 0.1% trifluoroacetic acid (TFA)/50% ACN, and 0.1% TFA/70% ACN, followed by concentration using a vacuum centrifuge.

The peptides were resuspended in 0.1% TFA and analyzed using an Easy-nLC1200 (Thermo Fisher Scientific) connected online to an Orbitrap Fusion LUMOS (Thermo Fisher Scientific) with a nanoelectrospray ion source (Thermo Fisher Scientific). Peptide samples were loaded onto a C18 analytical column (IonOpticks, Aurora Series Emitter Column, AUR2-25075C18A 25 cm × 75 μm 1.6 μm FSC C18 with nanoZero fitting) and separated using a 150-min gradient (solvent A, 0.1% FA; solvent B, 80% ACN/0.1% FA). For data-dependent acquisition of MS/MS spectra, the most intense ions (Cycle Time: 3 s) with charge state +2 to +7 were selected for fragmentation by higher-energy collisional dissociation (HCD) with a normalized collision energy of 30 and detected using Orbitrap. The resolution, duration of dynamic exclusion, and isolation window were set to 120,000, 15 sec, and 1.6 *m/z*, respectively.

The data were analyzed using a Sequest HT search program in Proteome Discoverer 2.4 (Thermo Fisher Scientific). MS/MS spectra were searched against the SwissProt-reviewed Homo sapiens reference proteome (UniProt v2017-10-25). Intensity-based label-free protein quantification was performed using a Precursor Ions Quantifier node in Proteome Discoverer 2.4. The mass tolerances for the precursor and fragment were 10 ppm and 0.02 Da, respectively. Maximum missed cleavage sites of trypsin were set to 2. Oxidation (Met), biotinylation (Lys), deamidation (Gln, Asn), and acetylation (protein N terminus) were selected as variable modifications. Peptide identification was filtered at FDR < 0.01.

### Quantification and statistical analysis

Statistical significance was determined by Fisher’s exact test, unpaired two-tailed t-test, or one-way ANOVA with post hoc test, testing for normal distribution and equal variance, unless otherwise indicated. Sample sizes, experimental replicates, and specific statistical tests used are described in the figure legends. Data were presented as means using GraphPad Prism 9 software. Error bars represent SD.

## Acknowledgements

We thank the FACS Core and the Microscopy Core Laboratories at the Institute of Medical Science, The University of Tokyo. This work was supported by Grant-in-Aid for Scientific Research (B) (22H03100, SG), Grant-in-Aid for Scientific Research on Innovative Areas (Research in a proposed research area) (21H00274, SG), Fostering Joint International Research (B) (22KK0127, SG), JSPS KAKENHI Grant Numbers JP22K16319 (KY), JP24K19216 (KY), JP23H05479 (YS), AMED under Grant Number (JP23ama221223, KY), research grant from the Japanese Society of Hematology (SG), the Tokyo Biochemical Research Foundation (KY), the Uehara Memorial Foundation (KY), and the Mochida Memorial Foundation for Medical and Pharmaceutical Research (KY). In addition to the aforementioned, we express our gratitude to all researchers who contribute to the study of MLOs and the development of biological tools related to our project. The achievements we have obtained are directly derived from their valuable findings, and we sincerely appreciate their efforts.

## Declaration of Interests

The authors declare that they have no conflict of interest.

## Author contributions

K.Y. designed and performed experiments, analyzed and interpreted the data, and wrote the manuscript. S.S. and Y. H. assisted in the experiments. T.T. and Y. S. performed LC/MS and analyzed these data. E.V. provided the BaEVRless plasmid and supervised the CRISPR/Cas9 knock-in methodology. S.G. conceived the project, designed the experiments, interpreted the data, and wrote the manuscript.

---

**Figure EV1 (related to Figure 1).**
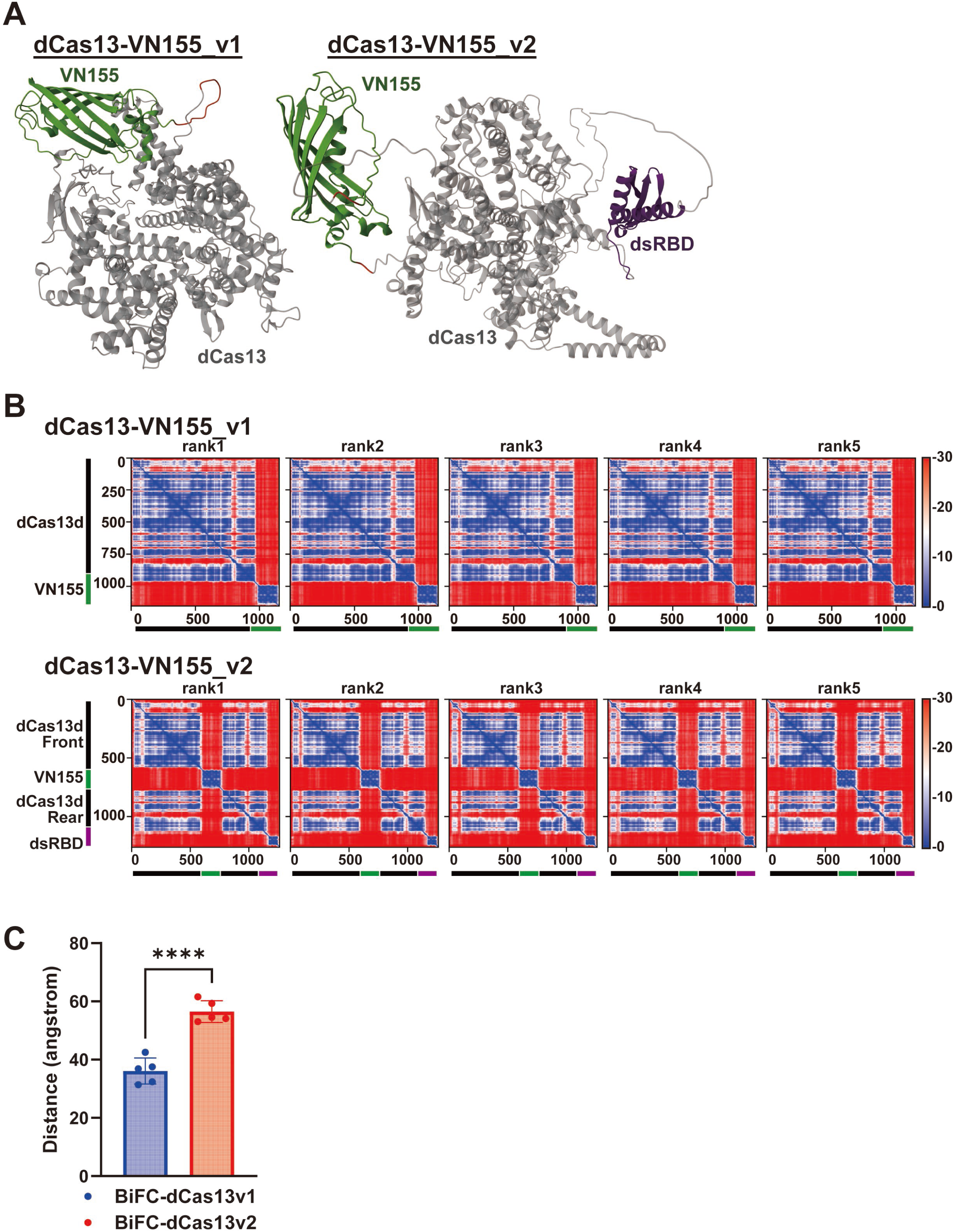
Structural features of BiFC-dCas13_v1 and BiFC-dCas13_v2. A. 3D protein structures of BiFC-dCas13d_v1 and BiFC-dCas13d_v2 predicted by AlphaFold. Structures illuminated in gray, red, green, and purple indicate dCas13d, flexible linker, VN155, and dsRBD, respectively. B. Solidity of the distance between each domain in dCas13-VN155_v1 (upper panels) and dCas13-VN155_v2 (lower panels) were predicted from amino acid sequences by AlphaFold. Five different models and their predictions are shown. C. Molecular distance between dCas13d and VN155 in prediction models of version 1 and version 2 are shown as individual data points and means +- SDs. For each dCas13d-VN155 version 1 and version 2, 5 prediction models were obtained from AlphaFold. Based on these models, center position of dCas13d and VN155 was determined and distance between each center positions were calculated through MolStar analysis. ****p<0.0001 (unpaired t test). N=5 for each group. See also Appendix Figure S1A.

**Figure EV2 (related to Figure 2).**
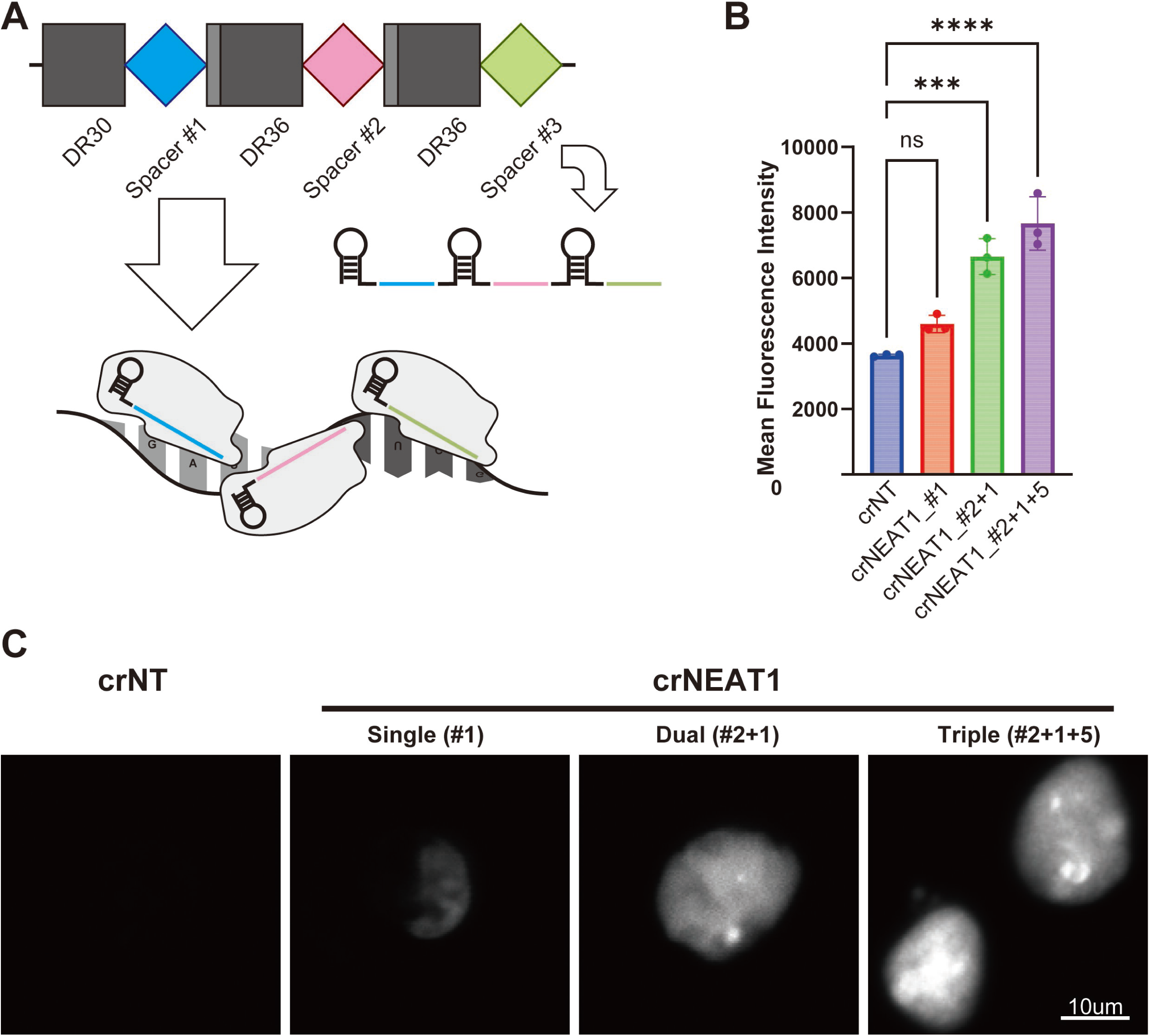
Triple crNEAT1 shows the highest signal intensity in the BiFC-dCas13 system. A. Schematics of triple-crRNA, and CRISPR/ Cas13 proteins guided to target transcripts. For triple-crRNA, 3 crRNAs targeting different position of same transcripts are tandemly placed. Each spacer sequences are followed by direct repeat (DR30 for first and DR36 for middle and last crRNA) and are processed into mature crRNAs. B. Live cell imaging of the BiFC-dCas13d_v2 systems combined with crNT, single-crNEAT1, dual-crNEAT1, and triple-crNEAT1. 293T cells were transfected with BiFC-dCas13d_v2 components and crRNAs. mVenus fluorescence images were acquired 2 days after transfection using the Nikon-A1 confocal microscope imaging system. Representative Venus fluorescence images of crNT (left), single-crNEAT1 (middle left), dual-crNEAT1 (middle right), and triple-crNEAT1 (right) are shown. Scale bars: 10 μm. C. Mean fluorescent intensity of the BiFC-dCas13d_v2 systems with the indicated crRNAs are shown as mean +- SDs. 293T cells were transfected with version 2 BiFC-dCas13d components and crNT, or three different types of crNEAT1. mVenus fluorescent signal is measured two days after transfection by FACS. ***p<0.001; ****p<0.0001 (one-way ANOVA with Tukey-Kramer’s post-hoc test). N=3 for each group.

**Figure EV3 (related to Figure 3).**
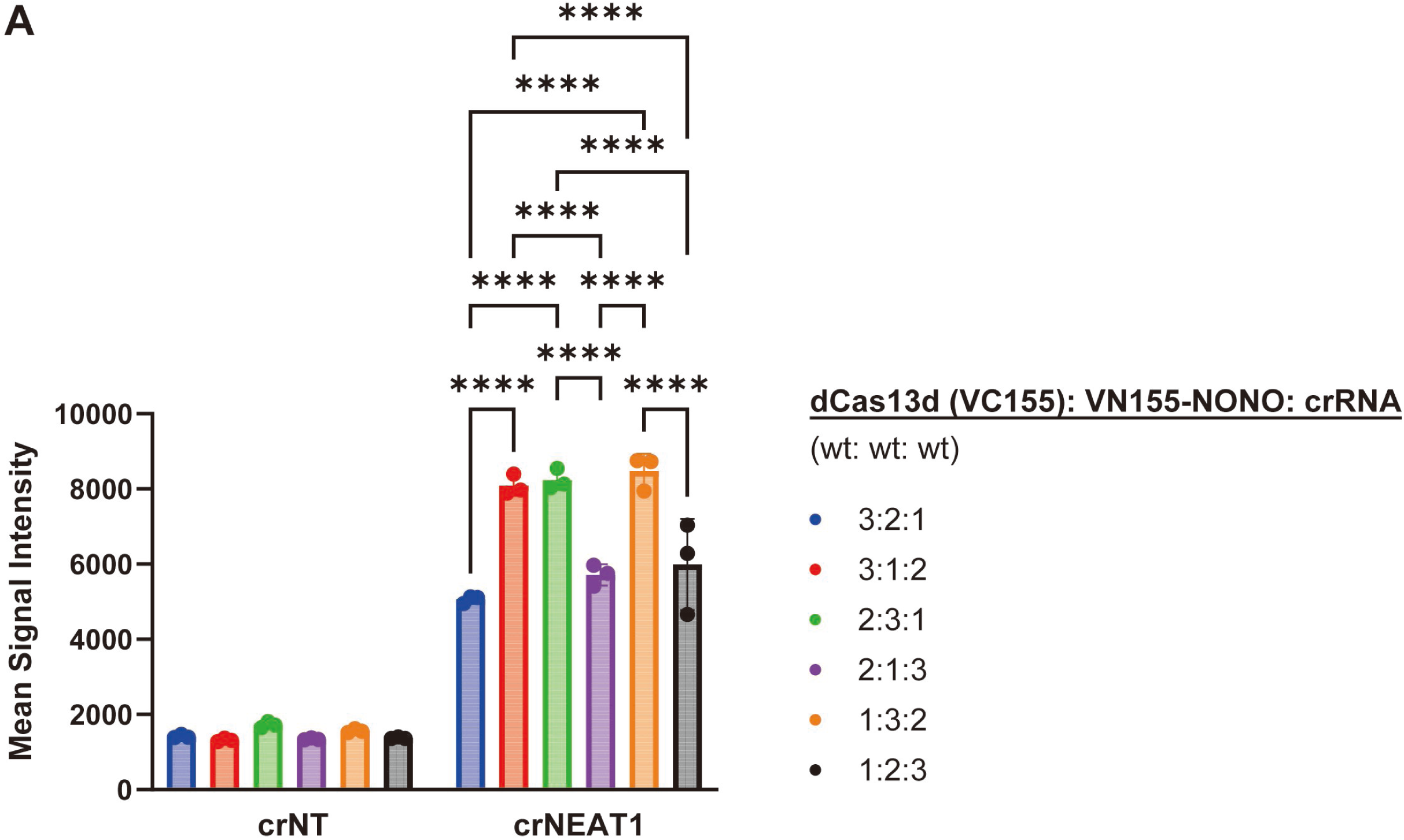
Stoichiometric verification of BiFC-dCas13d_3 systems. A. Mean fluorescent intensity of stoichiometrically varied BiFC-dCas13d_3 systems are shown as single data points and mean +- SD. 293T cells were transfected with BiFC-dCas13d_3 components at different molecular amounts as indicated. mVenus fluorescence signal is measured 2 days after transfection by FACS. ****p<0.0001 (one-way ANOVA with Tukey-Kramer’s post-hoc test). N=3 for each group.

**Figure EV4 (related to Figure 6).**
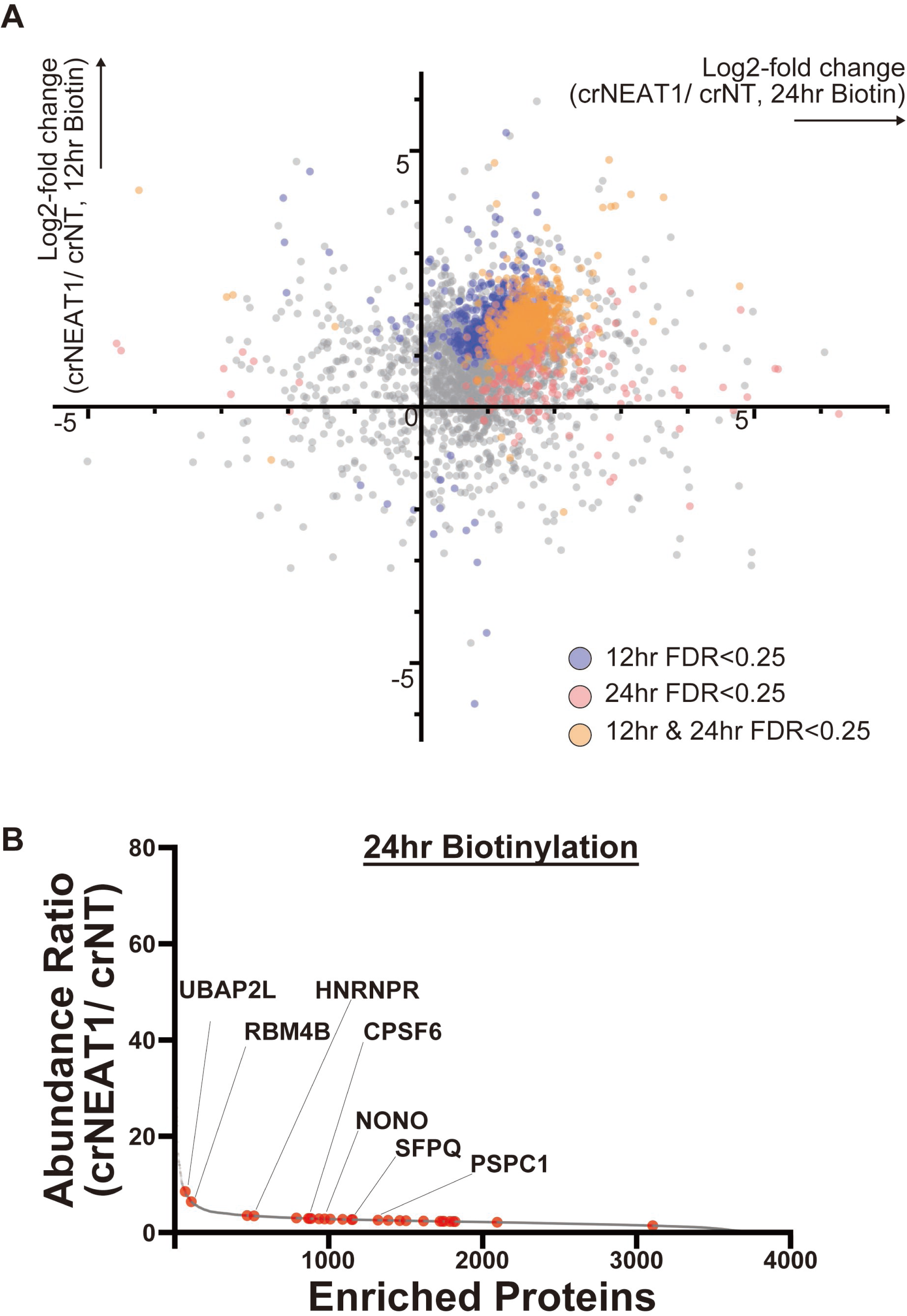
LC-MS/MS analysis of biotinylated proteins induced by Split-TurboID dCas13 system. A. Scatter plot showing log2-fold change (crNEAT1/ crNT, 24 hours biotinylation) versus log2-fold change (crNEAT1/ crNT, 12 hours biotinylation) for each protein. Blue, red, and orange dots indicate proteins showing statistical significance (FDR<0.25) in 12 hours biotinylation, 24 hours biotinylation, and both. B. Fold change (crNEAT1/ crNT) in protein abundance. Immunoprecipitated proteins after 24 hours of biotinylation induced by split-TurboID-dCas13d systems were subjected to LCMS/MS analysis. Red dots indicate proteins known as paraspeckle components. N=2 for each group.

**Figure EV5.**
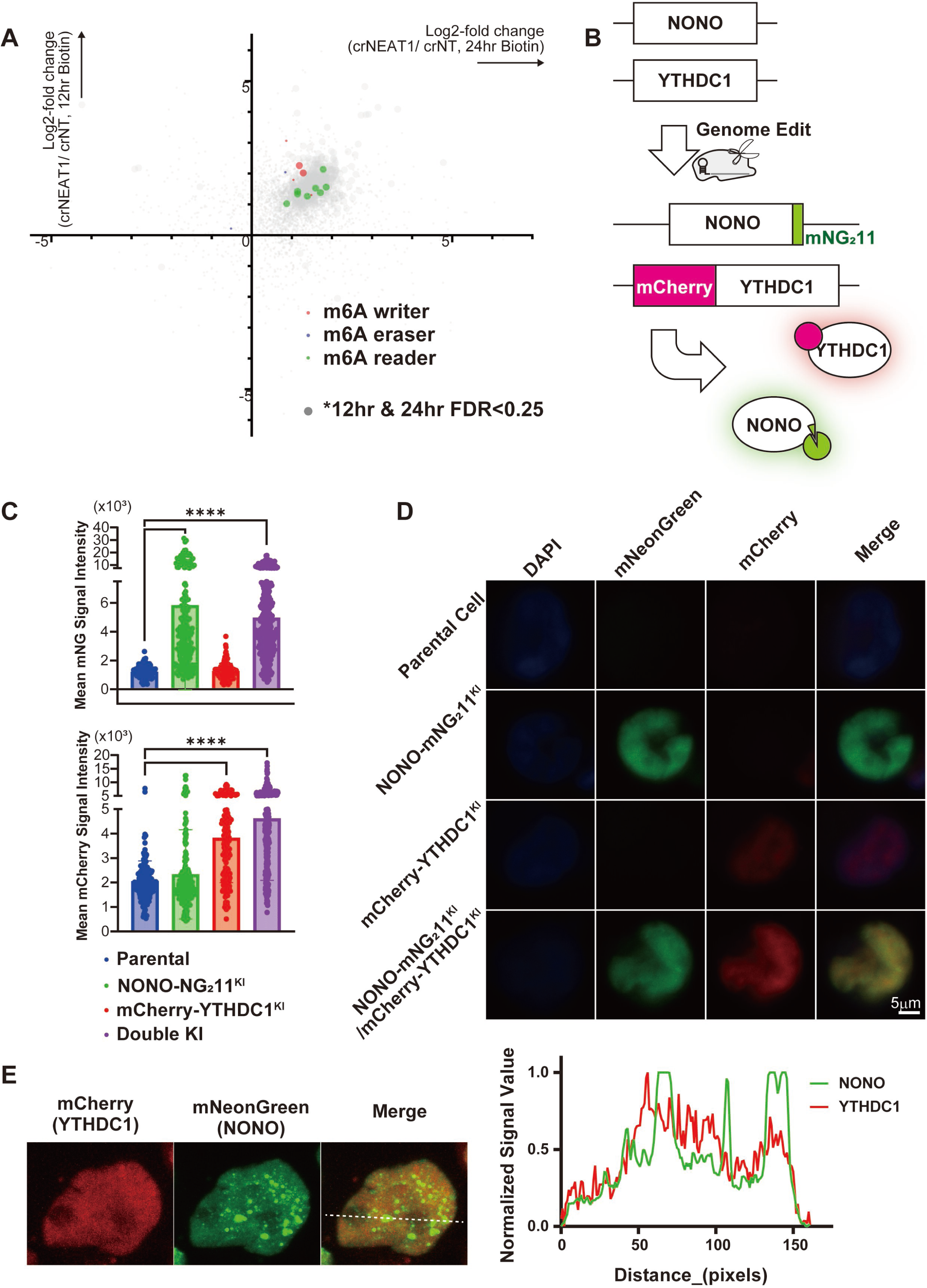
YTHDC1, m6A reader protein, interacts with NONO. A. Scatter plot showing log2-fold change (crNEAT1/ crNT, 24 hours biotinylation) versus log2-fold change (crNEAT1/ crNT, 12 hours biotinylation) for each protein related to Figure 7F. Red, blue, and green dots indicate m6A RNA regulatory proteins, writer, eraser, and reader respectively. Large dots indicate proteins that are significantly enriched by crNEAT1 (FDR<0.25) both at 12 and 24 hours. B. Schematic of gene editing in K562 cells. K562 cells expressing mNG3A (1-10) were subjected to genome editing to generate NONO-mNeonGreen (mNG) and mCherry-YTHDC1 knockin cells. mNG11 and mCherry protein coding DNA sequences were introduced directly into the C-terminus of NONO and N-terminus of YTHDC1, respectively, using CRISPR/Cas9. C. Mean signal intensities of mNG (left) and mCherry (right) in mNG11, mCherry, and dual knockin cells are shown as individual data points and mean +-SEMs. Gene-edited K562 cells were seeded on glass bottom dishes one day prior to observation. Fluorescence images were acquired using the Nikon A-1 microscope imaging system. Signal intensities were measured from 174, 203, 168, and 322 cells, respectively. ****P < 0.0001 (one-way ANOVA with Dunnett post hoc test). D. Representative images of parental, and mNG11/ mCherry knockin cells are shown. Scale bars: 5 μm. E. Representative live cell image of mNG11, mCherry, and dual knockin K562 cells (left). Cells were seeded on glass bottom dishes one day prior to observation. Fluorescence images were acquired using the Olympus IX73 equipped with a spinning disk confocal unit CSU-W1. Representative images are shown. Scale bars: 10μm. Plot profile analysis of signal intensity after min-max normalization (right). Signal intensities of mCherry (red) and mNeonGreen (green) were measured on the dotted lines shown in Figure EV5E.

